# Development of the Brief Personal Values Inventory for sense of values in the Japanese general population

**DOI:** 10.1101/479337

**Authors:** Sachiyo Ozawa, Yudai Iijima, Shuntaro Ando, Naohiro Okada, Tomoko Kawashima, Kazusa Ohta, Syudo Yamasaki, Kiyoto Kasai, Atsushi Nishida, Hironori Nakatani, Shinsuke Koike

## Abstract

The Portrait Values Questionnaire (PVQ) is a widely used questionnaire for assessing sense of values; however, it has limitations, especially for children. Therefore, the present study aims to develop a questionnaire for sense of values, called the Brief Personal Values Inventory (BPVI), consisting of simple questions and a smaller number of items compared to the PVQ. We first created 12 items for the BPVI and then tested their validity and reliability in 167 Japanese general population (81 males, mean age (SD) [range]: 23.4 (8.2) [15-57] years). Each of these items was correlated with one or more values in the PVQ-57. The BPVI items covered all higher-order values in Schwartz’s theory (Openness to Change, Self-Enhancement, Conservation, and Self-Transcendence). In sum, the BPVI has an acceptable criterion-related validity and corresponds to higher-order values in Schwartz’s value theory. The BPVI is suitable for a reliable and direct comparison of sense of values between children and adults, which may be useful for elucidating the developmental pathway of personal sense of values.

## 1. Introduction

Our personal sense of values determines our decision making, attitudes, and behaviors [1–3]. Values are thereby involved in all essential aspects of our lives including building and maintaining social relationships, developing religiosity, and making career choices that relate to our identity [3]. These aspects influence our quality of life, and thus, defining and studying sense of values has been regarded as an important topic.

Schwartz and his colleagues have been the first to classify sense of values into a systematic theory [2,4–6]. They initially proposed a theory about 10 motivationally distinct values founded on one or more universal human requirements for existence [6]. These values were structurally organized in a circular continuum according to the motivational goals they represent. Compatible values were located close to each other, and conflicting values in the opposite direction, based on two types of motivational dimensions (Personal Focus and Social Focus; Growth Anxiety-Free and Self-Protection Anxiety-Avoidance). The distinct values were also categorized into four higher-order values: Openness to Change, Self-Enhancement, Conservation, and Self-Transcendence.

Furthermore, they refined the value theory into more narrowly defined 19 values and developed a self-report questionnaire to measure the 19 distinct values, called 57-item Portrait Values Questionnaire (PVQ-57) [7]. The PVQ-57 was cross-culturally examined in 15 samples from 10 countries and showed a good cross-cultural validity. It became the most refined and widely used questionnaire for sense of values.

Most studies on personal sense of values, however, have been conducted with adult participants partly because PVQs have limitations when targeting children in particular. For assessing personal sense of values in children, the Picture-Based Value Survey for Children has been established (PBVS-C) [8]. However, a sentence-based questionnaire with simple and common questions that can be effectively applied to children has not yet been established. A previous study used different versions of PVQs to compare personal sense of values between children and adults: the PBVS-C for children and the 21-item PVQ for their parents [9]. However, reliable and direct comparisons of personal sense of values between children and adults have never been performed.

Personal sense of values is known to be transmitted within a family [10–13], and it is important to develop a questionnaire capable of directly comparing the differences in personal sense of values between children and their parents. Therefore, the present study intends to develop the Brief Personal Values Inventory (BPVI) to assess sense of values. We first created 12 items with simple questions for the BPVI and then examined the inventory’s validity and reliability through correlations with the PVQ-57, which represents the most refined value theory.

## 2. Materials and Methods

### 2.1 Participants

A total of 167 participants (mean age (SD): 23.4 (8.2) years, range: 15–57, 81 males) took part in this study. Participants were recruited through various types of advertisements, e.g., on bulletin boards in several high schools, job recruitment websites, and introduction by other participants, without any information about the survey or trial to avoid influencing the results. All participants were administered an anonymous self-report questionnaire in a school room. For test-retest reliability, we also obtained responses to the BPVI from 47 general women (51.2 (4.2) years) in the interval of 8 months by post. Participants were recruited to this study between June 2017 and July 2017.

The study was approved by the Research Ethics Committee at the Office for Life Science Research Ethics and Safety, The University of Tokyo (Approval No. 17-99 and 17-197). If a study intends to obtain data only using questionnaires, the ethical review guideline in Japan requires informed consent from only participants who are aged 15 or more and have graduated from junior high school or have equivalent capabilities. In this study, we recruited from high schools, and under the way approved by the ethical review board in the University of Tokyo, all participants gave written informed consent before participation according to the ethical approval. All of the authors did not have any access to information that could identify individual participants during or after data collection.

### 2.2. Measurements

#### 2.2.1. The 57-item Portrait Values Questionnaire (PVQ-57)

The PVQ-57 is a self-report questionnaire for assessing sense of values based on Schwartz’s refined value theory (Table 1 in S1 Appendix) [7]. The items assess how similar a participant is to the portrayed person (e.g., “It is important to him to form his views independently”). Each of the 19 distinct values is scored as the sum of three items responded on a 6-point Likert scale from 1 (“not like me at all”) to 6 (“very much like me”). The 19 values consist of two types of motivational dimensions (Personal Focus and Social Focus; Growth Anxiety-Free and Self-Protection Anxiety-Avoidance) and higher-order values (Openness to Change, Self-Enhancement, Conservation, and Self-Transcendence) [7]. The internal consistencies for the 19 value scores in this study sample showed acceptable Cronbach’s alpha greater than or equal to .66, except for the values of Self-direction–Thought (SDT) (.53) and Humility (HU) (.47) (Table 2 in S1 Appendix). Therefore, we excluded the values of SDT and HU from further analysis and finally scored for the 17 remaining values.

**Table 1.** The descriptive statistics of the 17 values in the PVQ-57.

|  |  | Mean (SD) |
| --- | --- | --- |
| **Higher-order values** |  |  |
|  | Openness to Change | 12.38 (2.56) |
|  | Self-Enhancement | 9.34 (2.73) |
|  | Conservation | 10.91 (2.15) |
|  | Self-Transcendence | 11.13 (2.29) |
| **Values** |  |  |
| SDA | Self-direction Action | 13.04 (3.16) |
| ST | Stimulation | 10.23 (3.18) |
| HE | Hedonism | 13.86 (3.00) |
| AC | Achievement | 10.73 (3.23) |
| POD | Power Dominance | 8.14 (3.19) |
| POR | Power Resources | 9.07 (3.36) |
| FAC | Face | 11.04 (3.18) |
| SEP | Security Personal | 12.39 (2.99) |
| SES | Security Societal | 10.88 (3.23) |
| TR | Tradition | 8.83 (3.13) |
| COR | Conformity-Rules | 10.56 (3.34) |
| COI | Conformity-Interpersonal | 11.86 (3.23) |
| BEC | Benevolence-Care | 11.53 (2.96) |
| BED | Benevolence-Dependability | 11.37 (3.20) |
| UNN | Universalism-Nature | 9.64 (3.01) |
| UNC | Universalism-Concern | 11.24 (3.45) |
| UNT | Universalism-Tolerance | 12.10 (3.10) |

**Table 2.** The descriptive statistics of the 12 BPVI items.

| Item | Content |  | Mean (SD) | N (%) | Skewness | Kurtosis |
| --- | --- | --- | --- | --- | --- | --- |
| B | beliefs | To have your own beliefs and to act on those beliefs | 7.6 (2.1) | 18 (1.8%) | -0.86 | 0.46 |
| I | interest | To pursue your interests | 7.9 (2.0) | 17 (1.2%) | -1.10 | 2.02 |
| AC | challenging | To actively seek for challenges | 7.2 (2.4) | 8 (4.8%) | -0.74 | 0.22 |
| L | leisure | To enjoy your hobbies or leisurely activities | 8.6 (1.8) | 29 (17.4%) | -1.98 | 5.40 |
| SI | social influence | To have social influence | 5.0 (2.9) | 3 (1.8%) | -0.09 | -0.76 |
| FS | financial success | To be financially successful | 7.3 (2.3) | 15 (9.0%) | -1.03 | 1.04 |
| GS | graduating school | To graduate from a prestigious school | 6.1 (3.2) | 2 (1.2%) | -0.64 | -0.64 |
| PE | positive evaluation | To be positively evaluated by others | 7.0 (2.5) | 6 (3.6%) | -1.20 | 1.08 |
| SL | stable-lifestyle | To maintain a stable lifestyle | 8.1 (2.1) | 20 (12.0%) | -1.54 | 2.78 |
| AT | Avoid causing trouble | To avoid causing trouble to others | 7.7 (2.4) | 11 (6.6%) | -1.24 | 1.03 |
| C | care | To care for close others | 8.5 (1.8) | 37 (22.2%) | -1.76 | 3.93 |
| IS | improving society | To improve society | 5.6 (2.5) | 0 (0.0%) | -0.17 | -0.40 |

N indicates the responses of the highest importance.

#### 2.2.2. Brief Personal Values Inventory (BPVI)

The BPVI intends to provide a simple and easy assessment for sense of values that can be applied to broader participants, including children. First, based on Schwartz’s 10 basic values, we selected and revised the nine items from PVQ-57: avoid causing trouble (AT), positive evaluation (PE), beliefs (B), improving society (IS), social influence (SI), challenging (AC), care (C), stable-lifestyle (SL), and leisure (L). We did not include the item for the value of Tradition (TR) (e.g., “It is important to him to follow his family’s customs or the customs of a religion”) because TR has been reported to be correlated with religiosity [14], and over 70% of Japanese people have been found to not have faith in only a specific religion [15]. We also created three additional items reflecting financial success (FS), graduating school (GS), and interest (I), which could motivate one’s behavior especially from an early stage of life. Consequently, 12 items were developed for the BPVI. Each question asks how important do participants consider the item for themselves. Participants responded on an 11-point scale ranging from 0 (“not important at all”) to 10 (“very important”). In a pre-survey, we confirmed that all items in the questionnaire were familiar for 12-year-old children from the general population in Japan.

### 2.3. Statistical analysis

All analyses were conducted using R version 3.4.4 and IBM SPSS statistics version 22.0. After descriptive statistics were examined, the relationships between the BPVI scores and age, sex, or present position (high school student, college student, employment, or disemployment or housekeeper) were tested. Subsequently, the criterion-related validity of the BPVI items was evaluated by Pearson’s correlation coefficient with the 4 higher-order value scores and the 17 value scores in the PVQ-57. In the correlation analyses, the relative BPVI scores were also used. The false discovery rate (FDR) [16] at the level of .05 was used.

## Results

The PVQ-57 and BPVI scores are shown in Tables 1 and 2, respectively. The mean (SD) of the BPVI scores was 7.2 (1.0).

Younger participants were more likely to ascribe greater importance to the higher-order values of Self-Enhancement (*r* = -.33, *p* < .001): Achievement (AC, *r* = -.31, *p* < .001), Power Dominance (POD, *r* = -.29, *p* < .001), and Power Resources (POR, *r* = -.22, *p* < .01) in the PVQ-57 (Table 3). The correlations were also seen in the values of Hedonism (HE, *r* =-.22, *p* < .01), Face (FAC, *r* = -.29, *p* < .001), and Benevolence–Dependability (BED, *r* = -.26, *p* < .001). Males ascribed greater importance than females to the higher-order values of Self-Enhancement (*t* = 4.25, *p* < .001): the values of AC (*t* = 4.72, *p* < .001) and POD (*t* = 3.69, *p*< .001). A one-way analysis of variance (ANOVA) revealed the main effect of present position in the higher-order values of Self-Enhancement (*F* = 6.10, *p* < .01): the values of AC (*F* = 5.11, *p* < .01) and POD (*F* = 4.76, *p* < .01). The differences were also seen in the value of HE (*F* = 4.56, *p* = .01) and BED (*F* = 4.56, *p* < .01). A Bonferroni post-hoc test showed that high school or college students scored greater than employed people and disemployed people or housekeepers.

**Table 3.** The demographics of the 17 values in the PVQ-57.

|  |  | Age | Sex |  | Present position |  |
| --- | --- | --- | --- | --- | --- | --- |
|  |  | *r* | *t* | *Dif* | *F* | *Dif* |
| **Higher-order values** |  |  |  |  |  |  |
|  | Openness to Change | -.11 | 2.11 |  | 2.98 |  |
|  | Self-Enhancement | -.33* | 4.25* | m > f | 6.10* | h > e, de; c > de |
|  | Conservation | -.13 | 0.31 |  | 2.79 |  |
|  | Self-Transcendence | -.04 | 0.92 |  | 1.30 |  |
| **Values** |  |  |  |  |  |  |
|  | SDA | .00 | 2.42 |  | 0.52 |  |
|  | ST | -.05 | 1.72 |  | 2.02 |  |
|  | HE | -.22* | 1.04 |  | 3.87* | h > e, de |
|  | AC | -.31* | 4.72* | m > f | 5.11* | h, c > de |
|  | POD | -.29* | 3.69* | m > f | 4.76* | h, c > de |
|  | POR | -.22* | 2.44 |  | 2.60 |  |
|  | FAC | -.29* | 1.73 |  | 3.52 |  |
|  | SEP | -.08 | 0.47 |  | 0.60 |  |
|  | SES | -.17 | 1.88 |  | 1.20 |  |
|  | TR | .01 | -0.27 |  | 1.21 |  |
|  | COR | -.10 | -0.49 |  | 3.32 |  |
|  | COI | -.18 | 1.16 |  | 3.58 |  |
|  | BEC | -.03 | 0.17 |  | 1.18 |  |
|  | BED | -.26* | 0.48 |  | 4.56* | c > e |
|  | UNN | .07 | 0.54 |  | 0.61 |  |
|  | UNC | -.02 | 0.74 |  | 0.50 |  |
|  | UNT | -.14 | 1.09 |  | 3.32 |  |

Younger participants were more likely to ascribe greater importance to PE (*r* = -.34, *p* < .001), GS (*r* = -.29, *p* < .001), and SI (*r* = -.27, *p* < .001) in the BPVI (Table 4). Males scored greater than females in the item SI (*t* = 2.97, *p* < .01). ANOVA revealed the main effect of present position for the item GS (*F* = 8.97, *p* < .001). High school and college students scored greater than employed people and disemployed people or housekeepers.

**Table 4.** The demographics of the 12 BPVI items.

| Item | Age | Sex |  | Present position |  |
| --- | --- | --- | --- | --- | --- |
|  | *r* | *t* | *Dif* | *F* | *Dif* |
| B | .04 | 1.41 |  | 1.84 |  |
| I | .01 | -0.37 |  | 1.18 |  |
| AC | .16 | 0.06 |  | 2.42 |  |
| L | -.03 | -0.09 |  | 1.05 |  |
| SI | -.27* | 2.97* | m > f | 1.41 |  |
| FS | -.16 | 2.08 |  | 1.38 |  |
| GS | -.29* | 1.29 |  | 8.97* | h, c > e; c > de |
| PE | -.34* | 2.58 |  | 3.18 |  |
| SL | -.17 | -0.37 |  | 4.02 |  |
| AT | .12 | -1.67 |  | 1.37 |  |
| C | .08 | -2.14 |  | 1.51 |  |
| IS | .11 | 1.55 |  | 0.18 |  |

m: male, f: female, h: high school student, c: college student, e: employment, de: disemployment or housekeeper
\* FDR corrected $p < .05$ .

Intraclass correlation coefficients for the BPVI items showed moderate to substantial test-retest reliability (AT, *ICC_(1,1)_* = .48; PE, .69; B, .52; FS, .41; IS, .67; I, .69; SI, .47; AC, .52; C, .51; GS, .60; SL, .71; L, .42).

### 3.1. Criterion-related validity of the BPVI items

Items B, I, AC, and L were associated with Openness to Change (Table 5). Items B, I, and AC were also significant but weaker associated with Self-Transcendence, located adjacent to Openness to Change and included in the dimension of Growth Anxiety-Free. Of these, item B was most significantly associated with the value of Self-direction–Action (SDA, *r* = .41, *p* < .001, Table 5), and items I and AC were most significantly associated with the value of Stimulation (ST, *r* = .38, *p* < .001, and *r* = .55, *p* < .001, respectively). Item L was correlated only with the value of HE (*r* = .35, *p* < .001).

**Table 5.**
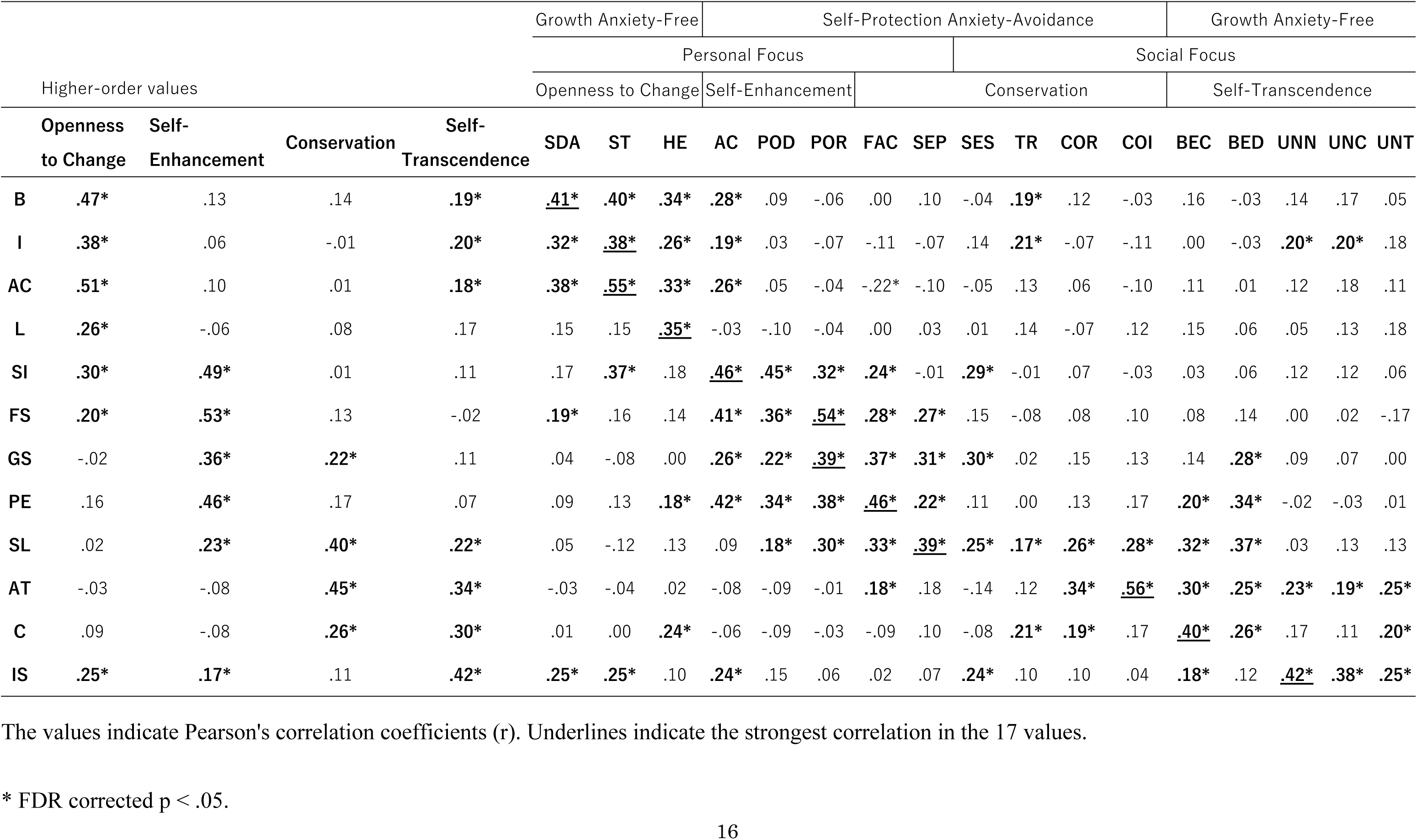
The correlations between the absolute BPVI scores and the 17 value scores in the PVQ-57.

Items SI, FS, GS, and PE were associated with Self-Enhancement. Similarly, the items SI and FS were also associated with Openness to Change, and the item GS with Conservation. For the correlation with the values, item SI was most significantly associated with AC (*r* = .46, *p* < .001), and items FS and GS were most significantly associated with the value of POR (*r* = .54, *p* < .001, and *r* = .39, *p* < .001, respectively). Item PE was most significantly associated with the value of FAC (Conservation, *r* = .46, *p* < .001).

Items SL and AT were associated with Conservation, and the strongest associated with the values of Security Personal (SEP, *r* = .39, *p* < .001) and Conformity-Interpersonal (COI, *r* = .56, *p* < .001), respectively. Item SL was also correlated with Self-Enhancement and Self-Transcendence, and AT with Self-Transcendence.

Items C and IS were correlated with Self-Transcendence, and the most significantly associated with the value of Benevolence–Care (BEC, *r* = .40, *p* < .001) and Universalism– Nature (UNN, r = .42, p < .001), respectively. Item C was also correlated with Conservation, and IS with Openness to Change and Self-Enhancement.

Compared with the absolute BPVI scores, positive correlations decreased and negative correlations increased when using the relative BPVI scores (Table 6). The strongest correlations were not different, except for the items SI and GS (Table 7).

**Table 6.**
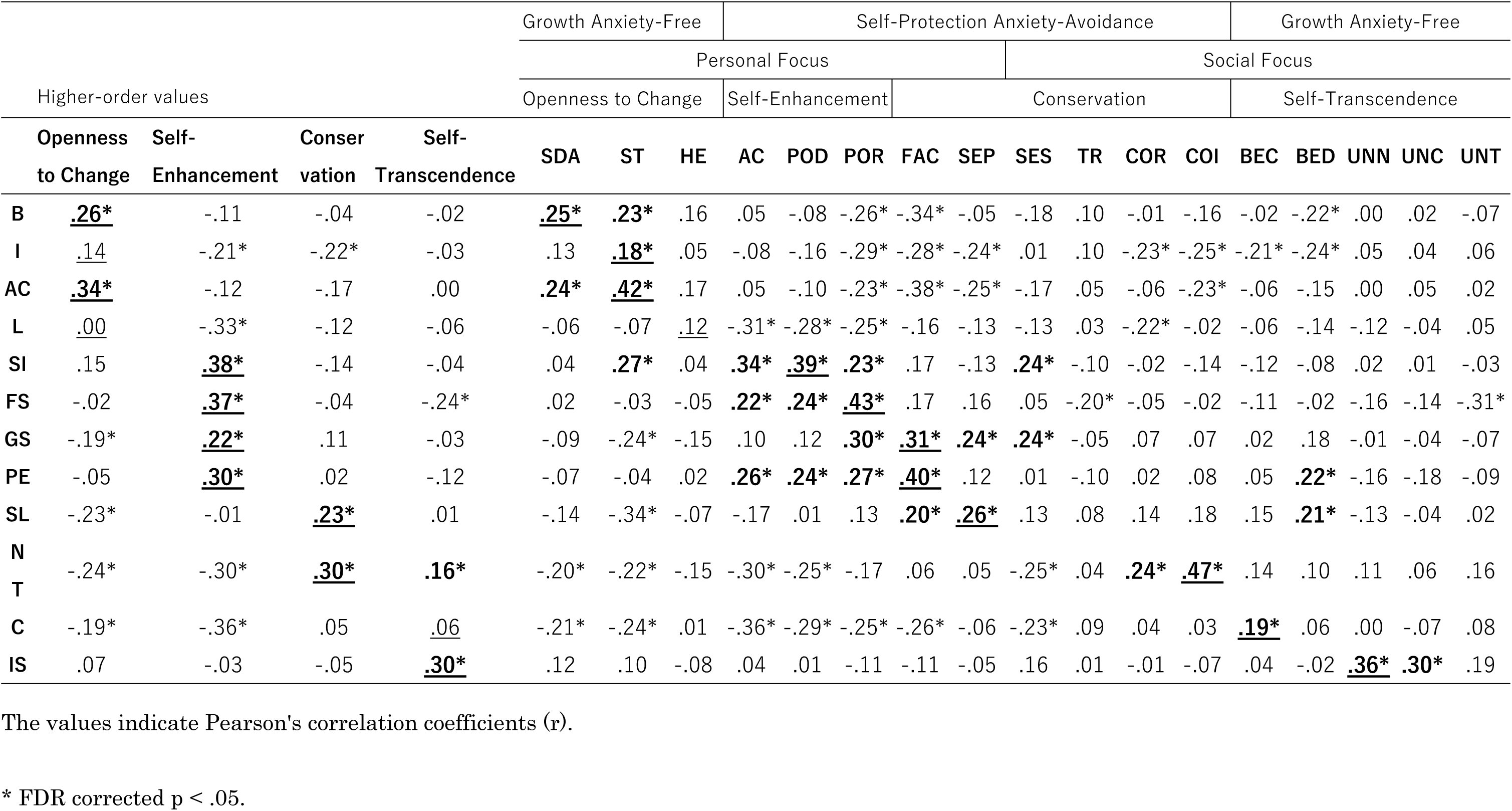
The correlations between the relative BPVI scores and the 17 value scores in the PVQ-57.

**Table 7.** The strongest correlations of the BPVI items with the values in the PVQ-57.

| 17 values<br>of the PVQ-57 | BPVI items |  |
| --- | --- | --- |
|  | Absolute | Relative |
| SDA | B | B |
| ST | I, AC | I, AC |
| HE | L | L |
| AC | SI |  |
| POD |  | SI |
| POR | FS, GS | FS |
| FAC | PE | GS, PE |
| SEP | SL | SL |
| SES |  |  |
| TR |  |  |
| COR |  |  |
| COI | AT | AT |
| BEC | C | C |
| BED |  |  |
| UNN | IS | IS |
| UNC |  |  |
| UNT |  |  |

## 4. Discussion

This study aimed to examine the validity and reliability of the BPVI, which assesses personal sense of values. The 12 items in the BPVI showed no ceiling or floor effects, and each item was correlated with one or more of the 4 higher order value and 17 values in the PVQ-57, indicating acceptable criterion-related validity. Four BPVI items were correlated with the higher-order value of Openness to Change, four items with that of Self-Enhancement, two items with that of Conservation, and two items with that of Self-Transcendence. Thus, the BPVI items were associated with a wide range of values in all higher-order values in Schwartz’s value theory.

Items B, I, AC, and L in the BPVI were associated with the higher-order value of Openness to Change in the Personal Focus and Growth Anxiety-Free dimensions of the PVQ-57. Item B “To have your own beliefs and to act on those beliefs” was most significantly associated with the similar concept of SDA (Freedom to determine one’s own action; Schwartz et al., 2012), and items I “To pursue your interests” and AC “To actively seek challenges” were most significantly associated with the value of ST (excitement, novelty, and challenge). Item L “To enjoy your hobbies or leisure activities” was unique in that it correlated only with the value of HE (pleasure and sensuous gratification), showing that it has good validity with HE. Items B, I, AC, and L were rarely correlated with the values of other higher-order values. Therefore, these items are conceptually similar to the higher-order value of Openness to Change in the PVQ-57.

The BPVI items SI, FS, GS, and PE were associated with the higher-order value of Self-Enhancement in the Personal Focus and Self-Protection Anxiety-Avoidance dimensions. Item SI “To have social influence” was most significantly associated with the value of AC (success according to social standards), and they both have the same characteristics. Item FS “To be financially successful” showed the highest correlation with the value of POR (power through control of materials and social resources). POR consists of items related to wealth (e.g., “It is important to him to have the power that money can bring”), and therefore, item FS is conceptually similar to POR. Item GS “To graduate from a prestigious school” was also most significantly associated with POR. Younger participants ascribed greater importance to item GS than older participants, whereas there was no difference in the value of POR. The meaning of item PE can be considered similar to that of either the items in the higher-order value of Self-Enhancement (e.g., “It is important to him that people recognize what he achieves”) or the value of FAC (e.g., “It is important to him that no one should ever shame him”). A confirmatory factor analysis (S1 Appendix) revealed that FAC cannot be included in the higher-order value of Conservation. The results suggest that item GS could be influenced by present student position. Items SI, FS, GS and PE were each correlated with all values in the higher-order value of Self-Enhancement, and they rarely correlated with the values in the opposite position.

Although items SL and AT were most significantly correlated with the values in the higher-order value of Conservation (FAC, SEP, and COI, respectively), these items were also broadly correlated with several values in more than one higher-order values. Item SL “To maintain a stable lifestyle” was correlated with all values in the higher-order value of Conservation, in addition to the conceptually similar adjacent values of POD (power through exercising control over people), POR, BEC (devotion to the welfare of ingroup members), and BED (being a reliable and trustworthy member of the ingroup). Thus, item SL corresponds well with the higher-order value of Conservation. On the other hand, item PE “To be positively evaluated by others” was associated with all values in the higher-order value of Self-Enhancement. Item AT “Avoid causing trouble to others” was correlated with all values in the higher-order value of Self-Transcendence. Conceptually, item AT is closest to the items in the value of COI (e.g., “It is important to him to avoid upsetting other people”). However, COI was associated with the values in the higher-order value of Self-Transcendence, rather than the values in Conservation, which may be a cultural characteristic of this study sample.

Items C and IS were most significantly associated with BEC and UNN (preservation of the natural environment) in the higher-order value of Self-Transcendence, as well as with the adjacent values of Conservation and Openness to Change, respectively. Item C “To care for close others” was conceptually similar to BEC, and the correlation was reliable. Item IS “To improve society” was conceptually similar to all values of universalism (UNN, UNC [commitment to equality, justice, and protection for all people], and UNT [acceptance and understanding of those who are different from oneself]). Therefore, items C and IS can be categorized into the higher-order value of Self-Transcendence.

This study has some limitations that should be mentioned here. First, since it is difficult for children to properly respond to the PVQ-57, in this study, we obtained responses from participants aged 15 years or more. Responses to several items and values varied by age, and thus, future studies need to investigate whether the BPVI could effectively assess sense of values in children. Second, relatively fewer items were correlated with the values in the higher-order values of Conservation and Self-Transcendence, and the dimension of Social Focus. Including additional items for them may help obtain a more detailed understanding of personal sense of values.

## Conclusions

This study tested the validity and reliability of the BPVI developed to assess personal sense of values. The results showed that each item in the BPVI corresponds to one of the higher-order values in Schwartz’s value theory. The BPVI consists of simple questions and can be applied to a wide variety of population, including children. Since the BPVI would enable a direct comparison between children and adults, it is a useful tool for assessing personal sense of values.

## Authors contribution

SO, YI, and SK wrote the draft manuscript and conducted statistical analyses. SA, NO, and SK made Japanese version of the PVQ-57 questionnaire. TK and KO contributed to recruitment and assessment from the participants. SY and AN made the BPVI questionnaire. All authors reviewed and have approved the final version of the manuscript.

## Supporting information

**S1 Appendix.**

**S1 Table. The items, 19 values, and higher order values in the PVQ-57 (Male Version).**

**S2 Table. The descriptive statistics and Cronbach's alpha of 19 values in the PVQ-57.**

**S1 Fig. Confirmatory factor analysis of the Japanese version of the PVQ-57 for higher-order values.**

**S3 Table. The correlations between the 19 values in the PVQ-57.**

## References

1. Rohan MJ. A rose by any name? The values construct. Personality and Social Psychology Review. 2000:4(3):255–277. doi: 10.1207/S15327957pspr0403_4

2. Schwartz SH. Universals in the Content and Structure of Values - Theoretical Advances and Empirical Tests in 20 Countries. Adv Exp Soc Psychol. 1992:25:1–65. doi: 10.1016/S0065-2601(08)60281-6

3. Sagiv L, Roccas S, Cieciuch J, Schwartz SH. Personal values in human life. Nat Hum Behav. 2017:1(9):630–639. doi: 10.1038/s41562-017-0185-3

4. Schwartz SH, Bilsky W. Toward a Universal Psychological Structure of Human-Values. Journal of Personality and Social Psychology. 1987:53(3):550–562. doi: 10.1037/0022-3514.53.3.550

5. Schwartz SH, Bilsky W. Toward a Theory of the Universal Content and Structure of Values - Extensions and Cross-Cultural Replications. Journal of Personality and Social Psychology. 1990:58(5):878–891. doi: 10.1037//0022-3514.58.5.878

6. Schwartz SH. Are There Universal Aspects in the Structure and Contents of Human-Values. J Soc Issues. 1994:50(4):19–45. doi: 10.1111/j.1540-4560.1994.tb01196.x

7. Schwartz SH, Cieciuch J, Vecchione M, Davidov E, Fischer R, Beierlein C, et al. Refining the Theory of Basic Individual Values. Journal of Personality and Social Psychology. 2012:103(4):663–688. doi: 10.1037/a0029393

8. Döring AK, Blauensteiner A, Aryus K, Drögekamp L, Bilsky W. Assessing values at an early age: The Picture-Based Value Survey for Children (PBVS-C). J Pers Assess. 2010:92(5):439–448. doi:10.1080/00223891.2010.497423

9. Döring AK, Makarova E, Herzog W, Bardi A. Parent-child value similarity in families with young children: The predictive power of prosocial educational goals. Br J Psychol. 2017:108(4):737–756. doi: 10.1111/bjop.12238

10. Whitbeck LB, Gecas V. Value Attributions and Value Transmission between Parents and Children. J Marriage Fam. 1988:50(3):829–840. doi: 10.2307/352651

11. Knafo A, Schwartz SH. Value socialization in families of Israeli-born and Soviet-born adolescents in Israel. J Cross Cult Psychol. 2001:32(2):213–228. doi: 10.1177/0022022101032002008

12. Knafo A, Schwartz SH. Parenting and adolescents’ accuracy in perceiving parental values. Child Dev. 2003:74(2):595–611. doi: 10.1111/1467-8624.7402018

13. Knafo A, Schwartz SH. Identity formation and parent-child value congruence in adolescence. Br J Dev Psychol. 2004:22:439–458. doi: 10.1348/0261510041552765

14. Saroglou V, Delpierre V, Dernelle R. Values and religiosity: a meta-analysis of studies using Schwartz’s model. Pers Indiv Differ. 2004:37(4):721–734. doi: 10.1016/j.paid.2003.10.005

15. Study of Japanese national character. The Institute of Statistical Mathematics. Retrieved 2018 Nov 7. Available from: http://www.ism.ac.jp/kokuminsei/table/data/html/ss3/3_1/3_1_all.htm

16. Benjamini Y, Hochberg Y. Controlling the False Discovery Rate - a Practical and Powerful Approach to Multiple Testing. J R Stat Soc Ser B-Methodol. 1995:57(1):289–300.

